# Plant-pollinator interactions in apple orchards from a production and conservation perspective

**DOI:** 10.1101/2023.11.20.567831

**Authors:** Anne-Christine Mupepele, Vivien von Königslöw, Anna-Maria Bleile, Felix Fornoff, Jochen Fründ, Alexandra-Maria Klein

## Abstract

In an agricultural landscape, production and conservation ideally go hand in hand. In a win-win scenario, conservation measures provide support for biodiversity and crop production, mediated by pollination for example. Hedges and flower strips are conservation measures that support pollinating insects, such as wild bees and hoverflies. They can be beneficial for crop pollination, but also harmful by dragging away pollinators from crops if flowering simultaneously. Here, we studied plant-pollinator interactions from two different perspectives. First we look at the apple-flower/production perspective investigating whether plant-pollinator networks in apple orchards differ with adjacent flower strips and hedges compared to isolated orchards. With help of the Bayes factor, we investigated similarity and conclude that there are no differences between pollination networks with or without adjacent flower strips and hedges. Second, we look at the pollinator/conservation perspective and analyse the impact of hedges and flower strips on pollinators and their interactions with plants before and after the apple bloom in April. We show that apple pollinators use more flower resources in flower strips and hedges across the whole season compared to isolated orchards. In orchards with flower strips and hedges interactions are more constant over time. We conclude that flower strips and hedges are beneficial for conservation of apple pollinators without being harmful for apple flower pollination being crucial for production.

## Introduction

Agricultural production relies on ecosystem services, such as pollination, which is essential for high yield quality and quantity in crops (Dainese *et al*., 2019; Garratt et al., 2014; Pardo and Borges, 2020; Palm et al., 2014; Tamburini et al., 2019). Depending on the crop species, insects, such as bees and hoverflies, are required for optimal pollination. Apple varieties generally depend to a certain degree on insect-mediated pollination (Pardo and Borges, 2020; Roquer-Beni *et al*., 2021). These are often honeybees purposefully managed, with hives placed next to orchards during apple bloom (Hung *et al*., 2019; Weekers et al., 2022). In addition to honeybees, the role of wild pollinators, such as wild bees and hoverflies has been recognized for many crops and for apple in particular (Garibaldi *et al*., 2011, 2013; Mallinger and Gratton, 2015; Rader *et al*., 2016; Page et al., 2021; Osterman et al., 2021b).

To support pollinators in agricultural landscapes, hedges and flower strips are politically promoted and hence planted and maintained in different places of Europe (Scheper *et al*., 2021; Garratt *et al*., 2017; Albrecht et al., 2020; Lowe et al., 2021; Eeraerts et al., 2021b). They are beneficial for pollinators as flower strips offer pollen and nectar from spring to late summer and with hedges playing an important role by offering early blooming floral resources (Hadrava *et al*., 2022; von Königslöw et al., 2022). Together these two pollinator conservation measures can support many pollinator species. Beside supporting pollinators, the additional flower offer in hedges and flower strips may also compete with simultaneously flowering crops (Holzschuh *et al*., 2016; Lundin et al., 2017; Osterman et al., 2021b). Such disservices are not in the interest of farming. Generally, farmers value pollination and are willing to support pollinators (Maas *et al*., 2021; Osterman et al., 2021a), but at best without disadvantages for production (KovácsHostyánszki *et al*., 2013; Mupepele et al., 2021).

Pollinators have species-specific nutritional requirements as they visit flowers of different plant species (Ruedenauer *et al*., 2019, 2020; Vaudo et al., 2015; Rodríguez-Gasol et al., 2020). A more diverse flower offer provided across the whole vegetation season generally results in a more diverse pollinator community (Glaum *et al*., 2021). Seasonal changes play a role as bees and hoverflies have specific flight periods which also vary in length, and plants are generally not flowering throughout the whole season. Plant-pollinator interactions thus change over the season (Balfour *et al*., 2018; CaraDonna et al., 2017; Bartomeus et al., 2013; von Königslöw *et al*., 2022).

Networks representing plant-pollinator interactions can improve our understanding of the effects of adjacent flowering conservation measures on pollinator-dependent crops (Rosa García and Minãrro, 2014; Bailes et al., 2015). Networks visualize species-specific flower visits of each pollinator species (Valido *et al*., 2019; Redhead et al., 2018) and reflect plant and pollinator relationships. In agricultural production with pollinator-dependent crops, they give insights to crop pollination with likely consequences to production.

While the influence of hedges, flower strips and other semi-natural habitats on pollinator diversity is well established (Scheper *et al*., 2015; Lowe et al., 2021), the influence on yield and plant-pollinator interactions in crop fields are less clear (Lowe *et al*., 2021; Albrecht et al., 2020). Some studies have found a benefit for yield, e.g. in strawberry (Grab *et al*., 2018), and others no relationship, e.g. in oilseed rape (Sutter *et al*., 2018). Apples are a frequently studied pollinator-dependent crop due to its high commercial importance in temperate climates (Pardo and Borges, 2020; Osterman et al., 2021b; Samnegård et al., 2019; Rosa García and Minãrro, 2014; Roquer-Beni et al., 2021; Garratt et al., 2021). But surprisingly few studies on interactions with pollinators and the resulting yield are available (Tamburini *et al*., 2019), and results are contradictory (Bishop *et al*., 2023; Campbell et al., 2017). Also the question, whether flower strips and hedges compete with apple flowers for pollinators, thus reducing pollination is so far less well known (but see Osterman *et al*., 2021b).

Pollinators need food resources beyond apple bloom and we assume that they are abundant in hedges and flower strips especially before and after apple bloom. While the available food offer for bees has been investigated in terms of floral abundance and diversity across the season (Balfour *et al*., 2018; Dainese et al., 2018; Glaum et al., 2021; Neumüller et al., 2021; von Königslöw *et al*., 2022), the changing interaction patterns of plants with pollinators and thus how different pollinator species use floral resources in orchard-adjacent flower strips and hedges before and after apple bloom is not well investigated.

In this study, we first analysed plant-pollinator interactions in apple orchards during apple bloom from an ‘apple-flower’/production perspective, and second plant-pollinator interactions in flower strips and hedges before and after apple bloom, taking the ‘apple-pollinator’/conservation perspective. We thus first investigate whether hedges and flower strips influence plant-pollinator networks in orchards during apple bloom hypothesizing that apple flowers are equally well pollinated independent of potentially competing adjacent conservation measures, such as flower strips and hedges. And second, whether apple-pollinating bees and hoverflies benefit from hedges and flower strips before and after apple bloom hypothesizing that apple pollinators benefit from hedges and flower strips across the whole season using a more diverse and abundant flower offer before and after apple bloom in orchards with adjacent flower strips and hedges. At the same time, we expect the number of plant-pollinator interactions to be more constant over time in orchards with adjacent flower strips and hedges.

## Methods

### Study area and design

Study sites were located in the south of Germany at the Lake Constance (Fig. 1a). Eighteen sites were chosen and categorised into four treatments: (i) apple orchards with an adjacent perennial flower strip planted in April 2018, (ii) apple orchard with an adjacent hedge at least 10 years old; (iii) apple orchards with an adjacent hedge and an additional flower strip (hedge herb layer) (iv) isolated orchards without any implemented conservation measures as controls (Fig. 1b; see von Königslöw *et al*., 2022, for flower strip species lists and further details, and Supplement A2). We used four to five replicates per treatment (see Fig. 1b).

**Figure 1:**
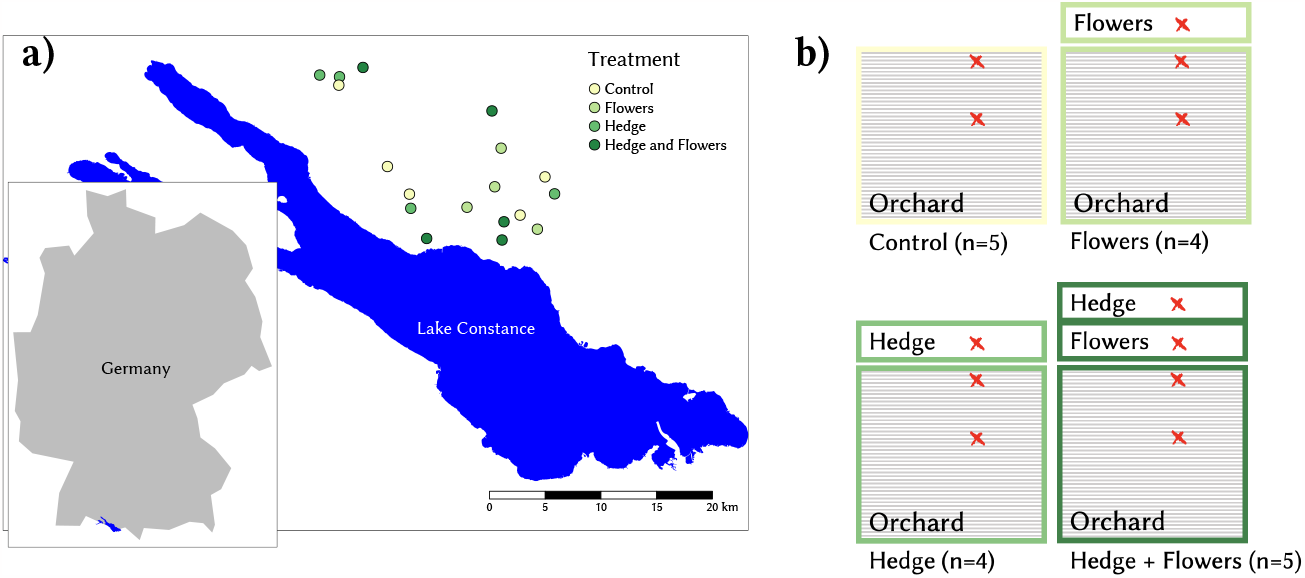
Study design showing the 18 study site locations at the Lake Constance (a) and the sample positions (red symbols) for the four different treatments (adjacent conservation measures) (b).

### Sampling method

Flower visits of bees (Apiformes) and hoverflies (Syrphidae) were sampled from March/April to August in 2018, 2019 and 2020. At each time an orchard study site was visited, one sample was taken from the inner apple orchard, one from the edge of the orchard and if present, one from the adjacent measures i.e. hedge, flower strip or one from both (Fig. 1b, red symbols for the sample location). Every sample consists of 15 minutes observations on three 1m2 rectangles (five minutes per rectangle, see von Königslöw et al. (2021) for further details). If possible pollinators and plants were identified to species level in the field, otherwise they were taken to the laboratory for further identifications. Sampling took place during good weather conditions meaning a temperature of at least 13°C, no precipitation and wind at less than 11m/s (on average 2.1m/s). Sampling effort varied between months and years, but study sites were sampled at least once per month, year and site. We subsumed most of the April samples under ‘Apple bloom’ to highlight the particularity of this month, while very few of the samples taken in the beginning of April, but before apple bloom were linked to the March samples and thus subsumed under ‘March’ in all figures. Each aggregation of samples was covering an approximate period of one month to avoid temporal aggregation on different scales (Schwarz *et al*., 2020).

### Statistical analysis

Flower visits of bees and hoverflies were visualized as plant-pollinator interactions in a bipartite plot and their properties were analysed with network indices. The analysis related to the apple-flower/production perspective is based on plant-pollinator interactions of all pollinators in orchards during apple bloom. This is a subset of the full data set only looking at samples taken during the apple bloom, discarding the other months, and only considering interactions taken in and at the edge of the orchards (Fig.1b, red symbols in the orchards). The analysis related to the apple-pollinator/conservation perspective is based on plant-pollinator interactions from all sample positions, i.e. in and at the edge of the orchards as wells as in flower strips and hedges across the whole season (Fig.1b, red symbols). For the apple-pollinator/conservation perspective only interactions with a pollinator that was at least once observed on an apple flower during apple bloom and thus assumed to be relevant for apple pollination was considered.

### Network index: Apple-flower/production perspective

For the apple-flower/production perspective, we calculated the network index species strength. The species strength is a species-level descriptor calculated as the sum of each species ‘dependencies’ (Eq.1; Bascompte et al., 2006; Dormann, 2011).

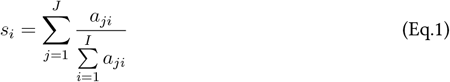

In Eq.1, *s*_*i*_ is the strength of the plant *i*, e.g. apple flowers. *a*_*ji*_ is the number of visits pollinator *j* pays to plant *i* (Bascompte *et al*., 2006).

The ‘apple species strength’ is thus reflecting the proportion of every pollinator species visiting apple flowers in relation to other orchard plants. The species strength is high if every pollinator species dedicates most of its visits to apples (in relation to other plant species in the network). We additionally analysed the abundance of pollinators visiting apple flowers, independent of the species identity of each pollinator and thus beyond a network.

### Network indices: Apple-pollinator/conservation perspective

For the apple-pollinator/conservation perspective, we calculated two indices: the pollinator generality and the effective number of partners. The network index ‘pollinator generality’ (Eq.2) is the number of plant species visited by a pollinator species and their even distribution on all plant species (Bersier *et al*., 2002; Dormann et al., 2008, 2009). Pollinator generality can be high if there are few pollinator specialists, but it is also an indicator for foraging choice and the presence of diverse and abundantly visited flowers (Doublet *et al*., 2022). If the pollinator species composition does not differ in terms of generalists and specialists, higher values stand for a high number of flowers and flower species visited by every pollinator, which means a more diverse food offer was used more evenly.

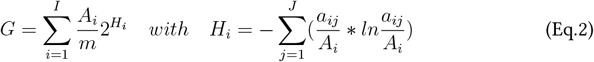

In Eq.2, *I* is the number of plant species (lower trophic level), *J* is the number of pollinator species, *m* is the total number of interactions, *a*_*ij*_ is the number of interactions between plant species *i* and pollinator species *j, A*_*i*_ is the total number of interactions of plant species *i* and *A*_*j*_ is the total number of interactions of pollinator species *j* (Bersier *et al*., 2002).

The second network index that we have used for the apple-pollinator perspective aims at identifying the stability and hence evenness of the number of interactions over time. The ‘effective number of partners’ index is the Shannon diversity to the power of e (Eq.3 Bersier et al., 2002; Jost, 2006; Dormann, 2011). .The index is high if the number of interactions from one site is evenly distributed across months (=‘partners’). The name of the index: ‘effective number of partners’ may be misleading in our context and we will call it ‘interaction constancy’, hereafter. We hypothesize that the index will be higher in orchards with adjacent conservation measures due to a more constant food offer and thus more constant interactions over time. The identity of plants and pollinators was not considered for this network index. The index is based on the same data than the pollinator generality index, hence apple-pollinators interacting with all plants in all sites, but considering site-month interactions with one event characterising any pollinator visiting any plant on a particular site in a particular month.

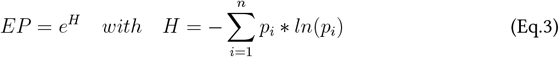

In Eq.3, *p*_*i*_ is the proportion of plant-pollinator interactions on a particular site per month *i* (Bersier *et al*., 2002).

The first two network indices (species strength (Eq.1) and pollinator generality (Eq.2)) were calculated by aggregating five random samples per month per site (drawn without replacement) to account for the different sampling effort. These five random subsamples were taken 100 times of the full dataset and aggregated. This resulted in 100 permutation rounds, each with one network index per month per site. The last index (interaction constancy) was standardized to sample effort by dividing the number of interactions per site and month by the number of samples taken in the respective site-month combination.

### Models and inference

The Bayes factor compares two competing models and can provide evidence for no effect, if the hypothesis is that there is no influence of a predictor variable in a linear model (Linde *et al*., 2023; Hartig and Barraquand, 2022). We have used the Bayes factor comparing two competing ANOVA models -one with treatment as predictor, the other without predictor variable-in combination with the response variables: apple species strengths, apple flower visits and interaction constancy. If the Bayes factor is below 1 the hypothesis in the denominator is favoured, which is in our case the intercept-only model (Linde *et al*., 2023). In this case we can conclude that the conservation measure has no influence on the respective plant-pollinator response variable.

Linear mixed models were used to identify whether the presence of adjacent conservation measures along the apple orchards and the season had an impact on the plant-pollinator network index ‘pollinator generality’ (Eq.4). As there were 100 network indices per site and month (from the repetitively taken random subsamples), we have estimated 100 models and computed their relevant parameters. Model characteristics were averaged across all 100 models and median and interquartile range are given. All analyses were realized in R version 4.2.3 (2023-03-15) (R Core Team, 2023), supported by the environment ‘RStudio’ Version 2023.03.1.

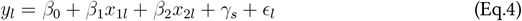

In Eq.4, *y*_*l*_ is the pollinator generality (*G*) per site and month; *x*_1_*l* and *x*_2_*l* are the two predictors treatment and month and *γ*_*s*_ is the random intercept on the *s*^*th*^ study site.

## Results

In total 5765 bees and 602 hoverflies were observed on flowers from 2018 to 2020. They were classified to 100 bee species and 22 hoverfly species which were recorded on 139 plant species. Honeybee was the most abundant pollinator species with 3918 specimen.

### Apple-flower perspective: Plant-pollinator networks during apple bloom

During apple bloom, apple flowers were visited predominantly by honeybees (see Fig. 2, orange interacting with purple). Wild bees and hoverflies contributed with 7% and 1% on average to apple flower visits (see Fig. 2, brown and blue). Beside apple trees, other plants were flowering in the herbaceous layer between trees in apple orchards. It was mostly dandelion (*Taraxacum officinale*), daisies (*Bellis perennis)*, bugle (*Ajuga reptans*) and ground ivy (*Glechoma hederacea*) (see Supplement A1 for networks with species identities). The pollinator community and their network links resembled between treatments regarding the proportion of honeybee, wild bees and hoverflies.

**Figure 2:**
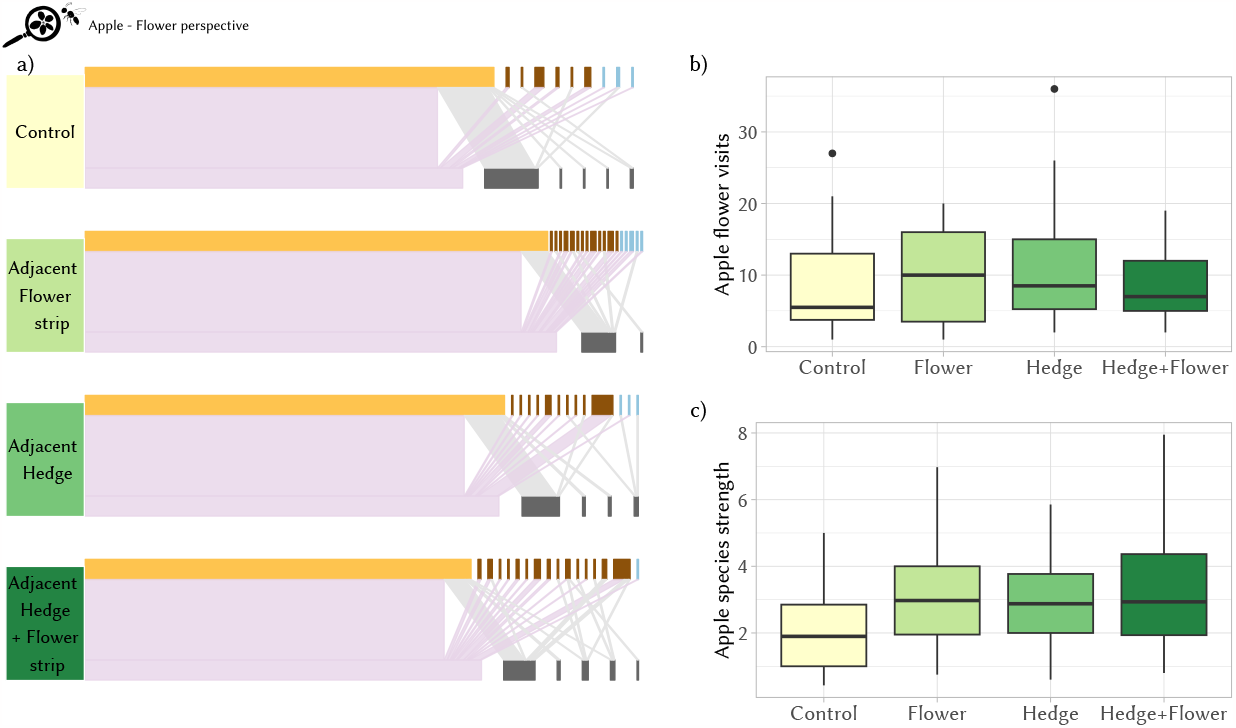
a) Plant-pollinator networks in apple orchards during apple bloom in the four treatments (green shades). Apple (purple) and other plants (grey) interact with honeybees (orange), wild bees (brown) and hoverflies (blue). Apple flower visits did not differ across treatments in terms of total number of pollinator visits (b) and visits in relation to other flowers in the network (network index: apple species strength, c)

Hedges and flower strips did not significantly influence apple flower visitation nor species strength (Table 1). Comparing the model with treatment as a predictor to an intercept-only model showed that the model without treatment as a predictor outperformed. In other words, we found no effect of hedges and/or flower strips on apple flower visitation and species strength. Apple flowers were visited equally often in all apple orchards and independent of adjacent hedges and flower strips (Table 1, Bayes factor = 0.22). Apple species strength, i.e. the apple flower visits per pollinator in relation to all other flower visits, was also independent adjacent hedges and flower strips (Table 1, Bayes factor =0.54).

**Table 1:** Results for the apple-flower perspective. Statistics for the species-strength model are averaged with median and interquartile range (IQR).

| Model | Analysis | Results |
| --- | --- | --- |
| log(Apple flower visits)<br>~ treatment | ANOVA | F = 1.39, p = 0.25, df = 3 |
| log(Apple flower visits)<br>~ treatment | Bayes Factor | BF = 0.22 |
| species strength ~ treatment | ANOVA | F = 0.99 (IQR = 0.78), p = 0.43 (IQR = 0.34), df = 3<br>average values from 100 models |
| species strength ~ treatment | Bayes Factor | BF = 0.48<br>average values from 100 models<br>in 94% of all subsampled models BF < 1 |

Apple flowers in orchards are equally well visited by pollinators across all treatments (Fig. 1). Hedges and flower strips lead to no benefit for apple pollination in terms of flower visits, but also to no disadvantage such as dragging away pollinators to adjacent hedges with simultaneously blooming flowers.

### Apple-pollinator perspective: Plant-pollinator networks across season

Apple pollinators interact not only with apple flowers, but a range of other plant species across the season (see Fig. 3a). Apple pollinators were mainly generalist species. Twenty-three of the 25 pollinator species occurred over more than one month and visited many different plant species (Fig. 3). The network index ‘pollinator generality’ reflects the number of plant species visited by pollinators and is weighted by the frequency each plant species was visited. Pollinator generality varied over the season and between the treatments (Fig. 3b, Table 2). Before apple bloom, orchards with hedges had on average a higher pollinator generality and were important (Fig. 3b). Most abundant in hedges were the wild bee species: *Andrena bicolor, Andrena haemorrhoa, Andrena stragulata, Colletes cunicularius, Osmia cornuta, Bombus lapidarius* and *Bombus terrestris*. They visited flowers of blackthorn (*Prunus spinosa*), European cornel (*Cornus mas*) and willow (*Salix sp*.) (Fig.3a). During apple bloom a generally lower apple-pollinator generality was observed. After apple bloom the control orchards remained low in terms of generality and clover (*Trifolium repens*) replaced apple flowers (*Malus sp*.) as the dominant species, but with reduced abundance (Fig. 3a). In flower strips the pollinator generality was significantly higher than in control orchards, most particularly in July (PostHoc results: Table 2). In orchards with flower strips and hedges, pollinators were more evenly distributed and visited more flowering species, thus demonstrating that they were offered a larger variety of food resources (Fig. 3b). This was also reflected in the network index ‘interaction constancy’ with a more evenly distributed number of interactions across the season in orchards with flower strips and hedges (Bayes factor = 1.7 pointed towards a likely effect, Fig. 4, Table 2).

**Table 2:** Results for the apple-pollinator perspective. Statistics for the pollinator-generality model are averaged with median and interquartile range (IQR).

| Model | Analysis | Results |
| --- | --- | --- |
| pollinator generality<br>~ season + treatment (Eq.4) | ANOVA | Season/Month: F=2.73 (IQR=1.44),<br>p=0.02 (IQR=0.065);<br>Treatment: F=3.55 (IQR=1.98), p<br>=0.039 (IQR=0.07)<br>average values from 100 models with<br>interquartile range (IQR) |
| pollinator generality<br>~ season + treatment (Eq.4) | PostHoc | Flower strip - Control: z=3.02, p=0.013;<br>July - Apple bloom: z=2.86, p=0.049 |
| interaction constancy ~ treatment | ANOVA | F = 3.06 , p=0.06, df = 3 |
| interaction constancy ~ treatment | Bayes Factor | BF = 1.7 |

**Figure 3:**
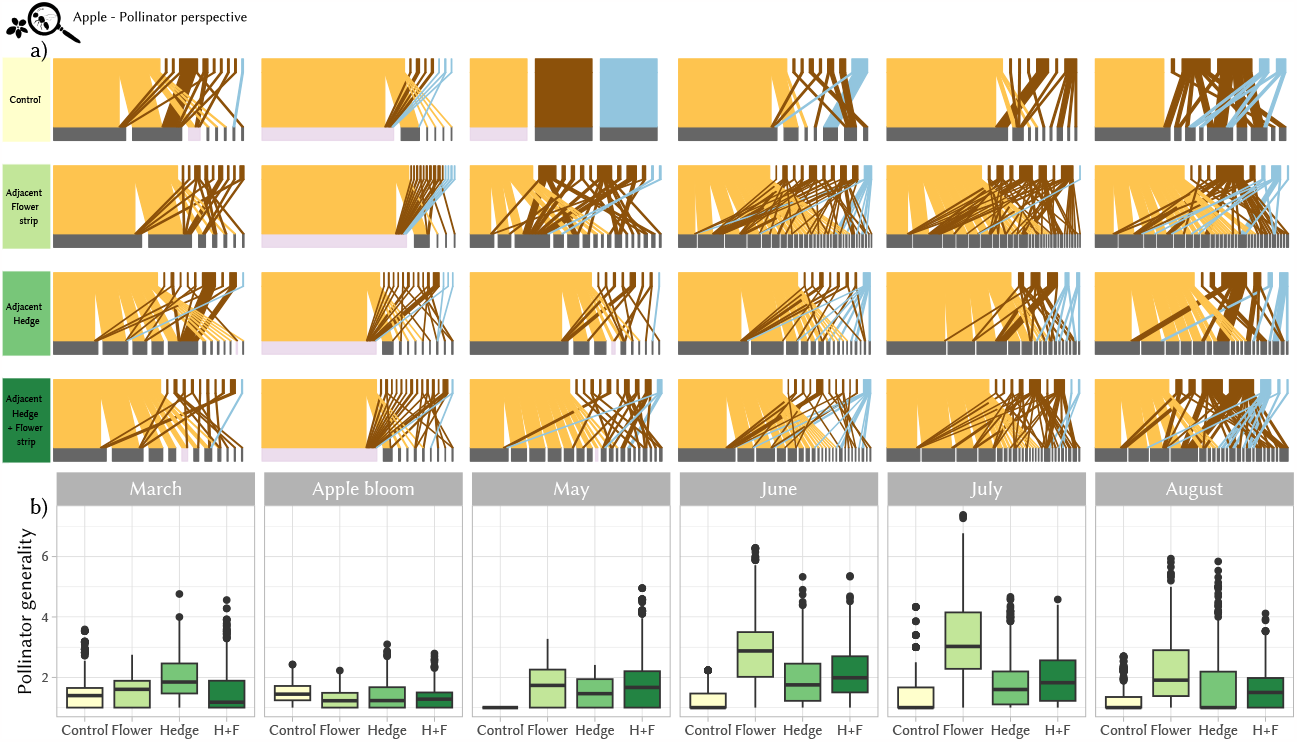
a) Plant - apple-pollinator networks in orchards and adjacent flower strips and hedges from March to August with Honeybees (orange), wild bees (brown), hoverflies (blue), apple flowers (purple) and other plants (grey). Conservation measures adjacent to orchards (treatments) are represented in different green shades. b) Pollinator generality differed between treatments (H+F = hedge and flower strip) and months.

**Figure 4:**
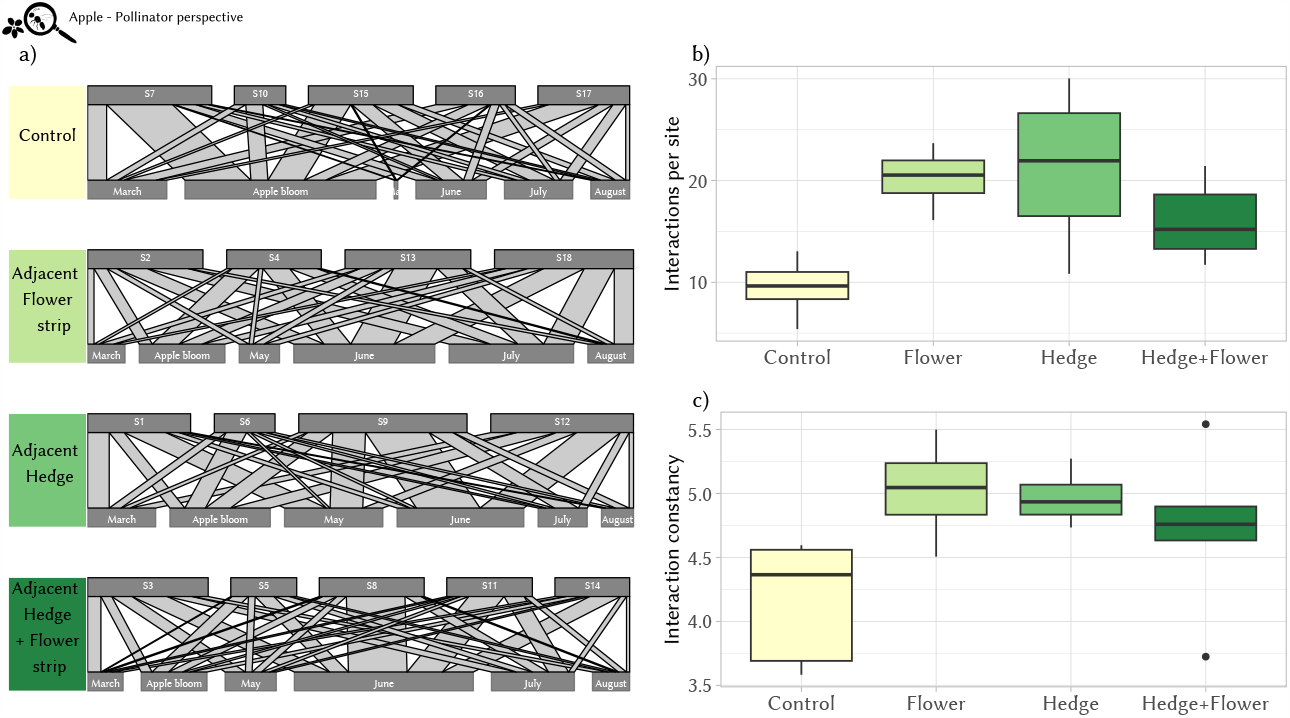
a) Site-month network with each interaction representing one plant-pollinator interaction per site and month. Boxplots with the number of interactions (b) and the interaction constancy (network index ‘effective partners’, c) demonstrate a higher and more even distribution of interactions across month in orchards with adjacent hedges, flower strips or both. Treatments (green) are orchards without adjacent conservation measure (‘Control’), with an adjacent flower strip (‘Flower’), with an adjacent hedge (‘Hedge’) or both (‘Hedge+Flower’).

## Discussion

Flower strips and hedges adjacent to apple orchards did not impact apple-flower visits and were likely to have no effect on crop pollination. However, flower strips and hedges were beneficial for apple pollinators, especially after the massflowering of apple in April. Apple-pollinating species, such as the bumblebee *Bombus terrestris* or the mason bee *Osmia bicornis*, benefit from a more constant flower offer, and here we showed that pollinators used the increased flower offer over the whole season. There was a more constant number of interactions across the whole flowering and flight period in orchards with hedges and flower strips compared to orchards without adjacent conservation measures.

### Apple-flower perspective

Apple production relies on pollination and thus farmers increasingly support pollinators by creating semi-natural habitats, such as flower strips or hedges. Flower strips and hedges can have an influence on the pollinator community in adjacent fields (Morandin and Kremen, 2013; Garratt *et al*., 2017; Albrecht et al., 2020; Lowe et al., 2021). In a beneficial sense, pollinators spill over to fields and pollinate crops (Holzschuh *et al*., 2012; Lowe et al., 2021; Ahrenfeldt *et al*., 2015). But plants blooming at the same time than apples might also draw away pollinators from the orchards and stand in competition with simultaneously flowering wild plants and crops (e.g. Kovács-Hostyánszki *et al*., 2013; Osterman et al., 2021b; Bishop et al., 2023). In our study, flower strips and hedges did not influence plant-pollinator interactions in apple orchards and the share every pollinator had to apple flowers in orchards was similar in all treatment no matter whether flower strips and hedges were present. Blackthorn (*Prunus spinosa*) and Willow (*Salix sp*.) flowering simultaneously to apple in hedges were not more attractive to pollinators. In cherry orchards, semi-natural habitat in the surrounding created a benefit for yield (Holzschuh *et al*., 2012), which we could not confirm for apple flower visitation. This was potentially due to a high amount of semi-natural habitats for all sites on a landscape scale that guaranteed a continuous resource availability, crucial for sustaining pollinator communities.

### Apple-pollinator perspective

Conservation of apple pollinators require flower resources. Flower strips and hedges provided such resources for apple pollinators, particularly before and after apple bloom. Looking at plant-pollinator networks across the whole season, the number of interactions of apple pollinators with non-apple flowers was constantly higher in orchards with adjacent flower strips and hedges in all month. Flower strips and hedges increased the diversity of plant resources used by pollinators. This confirms the conservation benefit of extending the availability of floral resources for apple pollinators (Carvell *et al*., 2022; Heller et al., 2019; von Königslöw et al., 2022).

Seasonality influences plant-pollinator interactions as not all pollinators and plant species are present throughout the season (CaraDonna *et al*., 2017; Bramon Mora et al., 2020). Some pollinator species have short flight periods, such as the European orchard bee, (*Osmia cornuta*) (compare Westrich, 2019). Other pollinators were present during the whole season, such as social species from the genus *Bombus* or *Lasioglossum* and those with several generations a year, e.g. *Andrena flavipes* or hoverflies. Regardless of phenological differences, pollinators could profit from the additional flower offer beyond the very short apple bloom. As apple pollinator species were mostly generalist species, they can use various flower species, if a diverse flower offer is present.

The interactions at our study orchards were dominated by honeybees. Honeybees are regularly used for apple pollination with honey bee hives hired during crop bloom. They are managed and fed if the flower offer is not sufficient. Nevertheless, not only honeybees, but most other apple pollinators, such as bumblebees and mason bees profited from flower rich habitat patches after mass-flowering (Riggi *et al*., 2021; Eeraerts et al., 2021a). Despite the dominance of honeybees, wild pollinators play an important role and can be more efficient for pollination services (Pardo and Borges, 2020; Földesi et al., 2016; Page et al., 2021). In our study both, honeybees and wild pollinators, were supported by orchard-adjacent flower strips and hedges.

Hoverflies are pollinators, but usually less efficient and often less abundant and less diverse, the latter was also the case in our study (Jauker *et al*., 2012; Rader et al., 2016; Pekas et al., 2020). Hoverflies in agricultural landscapes are often less sensitive to changes in land use and the spill over from semi-natural habitat to agricultural crops with hoverflies is more constant than for bees (Jauker *et al*., 2009). In terms of networks, we could not analyse differences between hoverfly and bee responses to flower strips and hedges, given the generally low number of hoverflies across all orchards.

### Beyond the orchard scale

Our study took place on an orchard-field scale and we have found that conservation measures did not influence plant-pollinator interactions during apple bloom, but had an impact after apple bloom. While mass-flowering of apple likely attracts and supports pollinators from the surrounding landscape, few pollinators were found inside the orchard after bloom with a lower flower offer in apple understories than in hedges and flower strips. (Riedinger *et al*., 2015; von Königslöw *et al*., 2022). This raises the question about the fate of apple pollinators in the orchards without any conservation measures after apple bloom and how or whether they could persist in the adjacent landscape. Apple pollinators have different abilities to cover distances between nest sites and foraging places (Hellwig *et al*., 2022). Hoverflies are not bound to any nesting habitat and can fly or drift across landscapes, especially if there are no high vegetation structures, such as hedges, which can pose barriers to hoverfly dispersal (Wratten *et al*., 2003). In contrast, bees are always bound to their nest site and require food resources within their flight range around the nest site, depending on the distance they can fly. This is generally not more than 500 m for smaller bees and up to 2.000-4.000 m for larger bees, but depends on the bee species and size (Zurbuchen *et al*., 2010; Földesi et al., 2016). If they nest in or close to orchards, apple pollinators must have found enough floral resources near the orchards, even when no conservation measures were implemented. Our isolated apple orchards (controls) were surrounded by other apple orchards and only very low proportions of forests (see von Königslöw *et al*., 2022). Nevertheless, small-scale semi-natural habitat patches like drainage ditches or slopes can provide noteworthy floral resources in the agricultural landscape matrix (Librán-Embid *et al*., 2021; von Königslöw et al., 2021). If we look at larger scales, the lack of semi-natural habitat has a negative impact on crop yield and plant-pollinator relationship (Garibaldi *et al*., 2011; Holzschuh et al., 2012; Földesi et al., 2016; Kleijn et al., 2015). Potentially the landscape scale could buffer the effect on the field scale for apple flower pollination.

### Conclusion

We have found that apple flower visits are not disadvantaged by conservation measures adjacent to orchards, and apple production does not stand in competition to hedges or flower strips. At the same time, apple pollinators, such as the early mining bee (*Andrena haemorrhoa*), profit from such conservation measures before and after apple bloom. For apple production and farming, it is thus favourable to implement conservation measures as there are no disadvantages for apple pollination, but benefits for apple pollinators after apple bloom. Conservation measures, such as flower strips and hedges, can likely help to stabilize unmanaged apple pollinator populations in an agricultural landscape.

## Acknowledgements

We thank the orchard owners and field assistants for their support during field work. The research project was funded by Bayer Crop Science. The first author was also supported by the Ministry Of Science, Research and the Arts Baden-Württemberg from August 2021 to April 2023.

## Supplement A1

**Figure A1:**
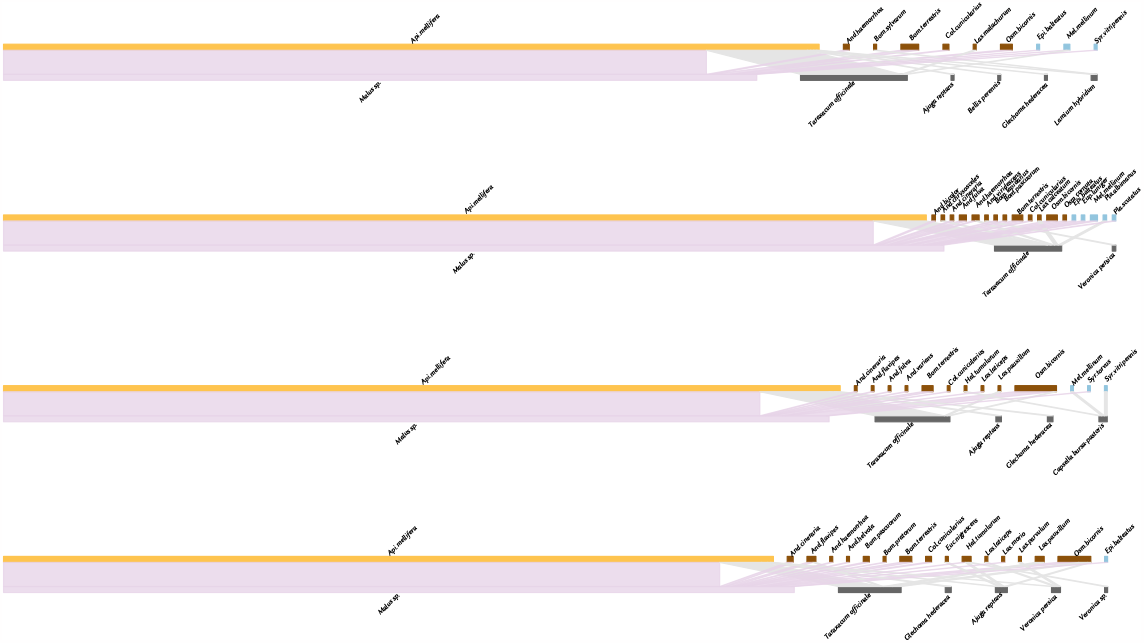
Supplementary figure with plant-pollinator networks in apple orchards during apple bloom in the control sites, sites with adjacent flower strips, with hedges and with hedges and flower strips (from top to bottom). Apple (purple) and other plants (grey) interact with honeybees (orange), wild bee (brown) and hoverfly species (blue).

### A2

zip file with R-Markdown document, html and data csv to reproduce the analysis.

